# Inferring biosynthetic and gene regulatory networks from *Artemisia annua* RNA sequencing data on a credit card-sized ARM computer

**DOI:** 10.1101/661058

**Authors:** Qiao Wen Tan, Marek Mutwil

## Abstract

0.

Prediction of gene function and gene regulatory networks is one of the most active topics in bioinformatics. The accumulation of publicly available gene expression data for hundreds of plant species, together with advances in bioinformatical methods and affordable computing, sets ingenuity as the major bottleneck in understanding gene function and regulation. Here, we show how a credit card-sized computer retailing for less than 50 USD can be used to rapidly predict gene function and infer regulatory networks from RNA sequencing data. To achieve this, we constructed a bioinformatical pipeline that downloads and allows quality-control of RNA sequencing data; and generates a gene co-expression network that can reveal enzymes and transcription factors participating and controlling a given biosynthetic pathway. We exemplify this by first identifying genes and transcription factors involved in the biosynthesis of secondary cell wall in the plant *Artemisia annua*, the main natural source of the anti-malarial drug artemisinin. Networks were then used to dissect the artemisinin biosynthesis pathway, which suggest potential transcription factors regulating artemisinin biosynthesis. We provide the source code of our pipeline and envision that the ubiquity of affordable computing, availability of biological data and increased bioinformatical training of biologists will transform the field of bioinformatics.

**Highlights:**

- Processing of large scale transcriptomic data with affordable single-board computers
- Transcription factors can be found in the same network as their targets
- Co-expression of transcription factors and genes in secondary cell wall biosynthesis
- Co-expression of transcription factors and genes involved in artemisinin biosynthesis

## 1. INTRODUCTION

To extract useful knowledge from rapidly accumulating genomic information, we are dependent on our capacity to correctly assign biological functions to gene products. Since gene products often have more than one function that may be carried out under different conditions, functional characterization of a gene can take years. This explains why, despite of decades of intensive research, only 12 % of genes in Arabidopsis thaliana (3382 out of 28392 genes) are functionally characterised as of 2018 [1].

To facilitate functional characterisation, bioinformatical gene function prediction can help experimentalists by (i) identifying novel genes relevant for the biological process studied and (ii) by suggesting relevant experimental approaches to dissect the function of an unknown gene.

For example, several studies used *in silico* predictions to identify genes involved in seed germination [2], plant viability [3], cyclic electron flow [4], cell division [5], shade avoidance [6], drought sensitivity and lateral root development [7]. Consequently, gene function prediction is one of the most intensive bioinformatics research efforts [1,8–10].

Apart from predicting gene function, a substantial effort in bioinformatics is given to identifying transcription regulatory networks, especially transcription factor (TF) - target associations. TFs regulate their target genes by binding to short DNA sequences called TF binding sites (TFBSs). The full set of regulatory interactions between a TF and its target genes forms a gene regulatory network (GRN). Since gene expression regulation is fundamental to all life, unraveling the GRN is pivotal to understand how different biological processes such as development, growth and stress responses are controlled.

GRNs are typically elucidated by yeast one-hybrid (Y1H), which tests for binding of a transcription factor to a promoter. Y1H was used to unravel the root-specific GRN in Arabidopsis [11], secondary cell wall synthesis [12], regulators for SHORTROOT-SCARECROW [13] and others. Further experimental methods include ChIP-chip or ChIP-Seq methods that determine TF binding to cis-elements *in vivo* [14], open chromatin profiling by DNAse I hypersensitivity [15], protein binding microarrays [15], DNA affinity purification assays [16] and others. While invaluable, the experimental methods to uncover GRNs have limitations generally inherent to methods used to elucidate gene function, as e.g. ChiP-Seq is labor intensive and does not allow a systematic dissection of genome-wide GRNs [17]. In these cases, computational predictions have proven highly efficient in supporting the elucidation of GRNs. For example, by integrating regulatory interactions from publicly available data, AtRegNet (https://agris-knowledgebase.org/, [18]) allows the visualisation of complex networks formed by TFs and their target genes. Other approaches take advantage of cross-species or cross-ecotype conservation of non-coding regulatory sequences in promoters [19–22]. These approaches can be further augmented by integrating gene expression data to identify transcriptionally co-regulated (co-expressed) genes, that are often functionally related (http://bioinformatics.psb.ugent.be/webtools/TF2Network/, [23–25]).

In general, the gene function and GRN prediction methods can be classified into three “generations”: (i) single, (ii) integrative and (iii) ensemble [24]. Regardless of the method, these approaches are based on the guilt-by-association principle, where genes with similar features are assumed to have the same function [10,26]. First generation methods use a single data type to predict gene function or GRN. For example, sequence similarity by BLAST can reveal proteins with a similar sequence, while analysis of gene expression can reveal co-expressed genes [10,23,24]. Second generation methods integrate multiple data types, thus increasing the coverage of the functional associations [10] and decreasing false positives [27–29]. Third□generation ensemble (also called community) methods integrate the predictions of many first□ and second□generation methods. Integration of 29 predicted GRNs in yeast and *E. coli* produced an ensemble GRN that outperformed the individual methods [30]. In plants, ensemble methods have been applied to predict subcellular localization of proteins (http://suba3.plantenergy.uwa.edu.au/, [31]), GRNs [32] and gene function [1].

While co-expression networks are not typically used to infer GRNs, we were able to successfully use these methods in our studies on secondary cell wall biosynthesis pathways [23,24,33,34]. This shows that GRNs can be elucidated by co-expression analyses for biological processes and pathways if they are under strong transcriptional control, such as secondary metabolism.

In this paper, we demonstrate how GRNs can be rapidly inferred on the Rock64, a device worth less than 50 USD, by taking advantage of the recent advances in affordable single board computers (https://store.pine64.org/?product=rock64-media-board-computer) and efficient RNA-seq quantification programs [35]. The Rock64 contains a quad-core ARM-based central processing unit, 4 GB RAM, numerous input/output ports (Figure 1A) and supports various Linux-based operating systems.

**Figure 1.**
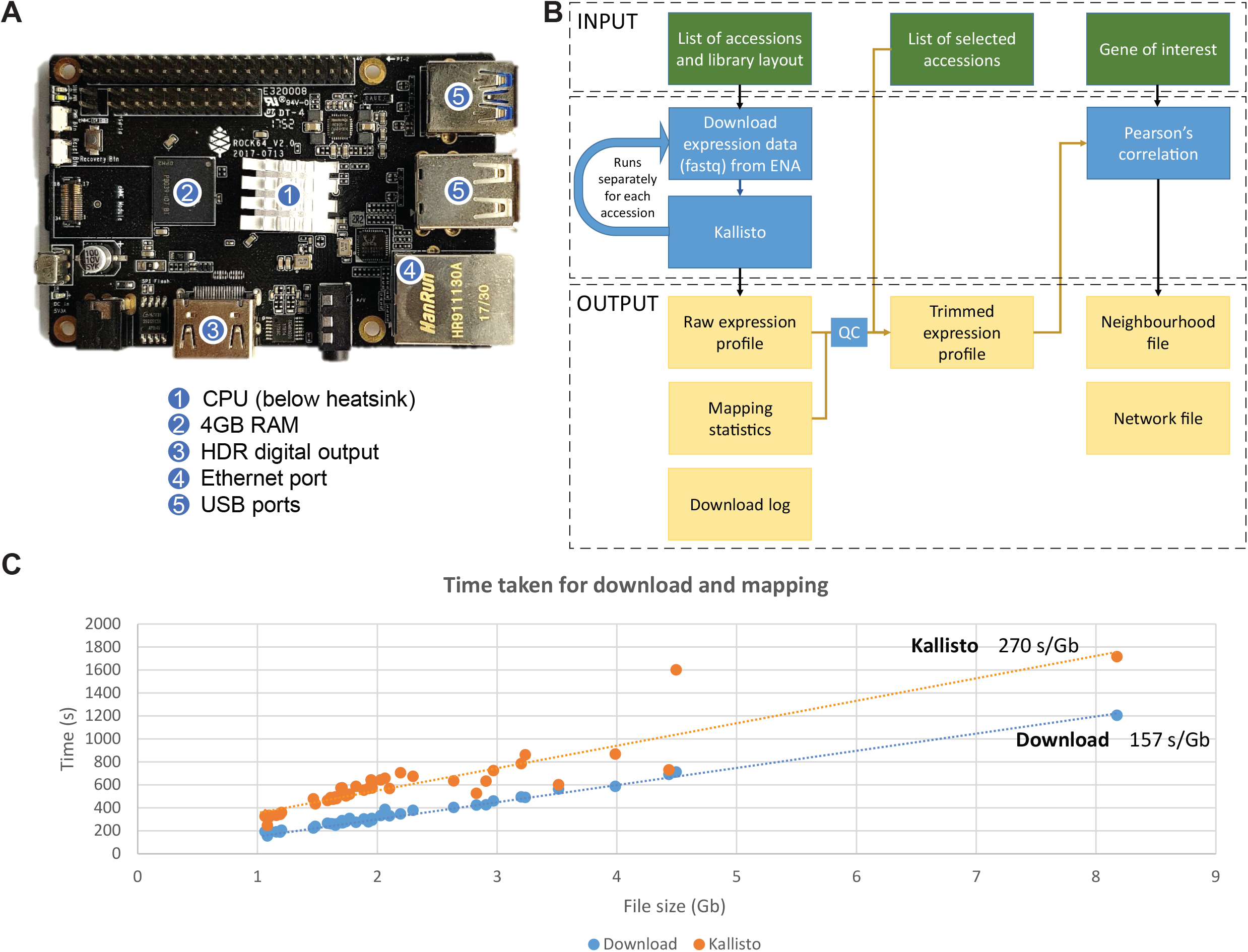
Establishing RNA sequencing and co-expression network construction pipeline on Rock64. A) Anatomy of Rock64 single board computer. The major components are 1. central processing unit (CPU), 2. 4GB RAM, 3. HDR digital output, 4. Ethernet port, 5. USB ports. B) Flowchart schematic for the pipeline used to download and process the RNA sequencing data and construct a co-expression network. C) Download and kallisto mapping times for the *Artemisia annua* data.

Next-generation sequencing technologies are currently outpacing Moore’s law, which reflects a trend observed in the computer industry that involves the doubling of computing power every two years [36,37]. The cost per genome has dropped from 100 million USD to 1000 USD in less than two decades (https://www.genome.gov/about-genomics/fact-sheets/DNA-Sequencing-Costs-Data). Similarly, the sequencing hardware has become more affordable and compact, changing from bulky, expensive devices weighing tens of kilograms and costing >100,000 USD, to portable sequencers weighing <100 grams and costing <1000 USD. The improvements in software needed to process sequencing data have followed similar trends. For example, kallisto and salmon, programs used to estimate gene expression are at least 1000 times faster and require less memory than the previous generation of corresponding software [35,38]. Consequently, RNA sequencing analyses that would take weeks on an expensive computer cluster can now be performed in a matter of days on a laptop. These developments prompted us to showcase how to use the Rock64 to process the available RNA sequencing data for *Artemisia annua*, to infer biosynthetic and GRN networks of secondary cell wall and artemisin biosynthesis.

## 2. MATERIAL AND METHODS

RNA sequencing experiments for *Artemisia annua* was downloaded as fastq files from European Nucleotide Archive (ENA, [39]) via Aspera v3.5.6.110647. To run Aspera on the ARM CPU of Rock64, we have used x86 CPU emulator ExaGear (https://eltechs.com/product/exagear-desktop/). Using Aspera is optional, but it provides fast download speeds. For paired-end experiments, only the file containing the first read, designated with “_1” was downloaded. The mapping of reads was done using kallisto v0.44.0 [35]. Kallisto index file was generated for *A. annua* (https://www.ncbi.nlm.nih.gov/assembly/GCA_003112345.1/, [40]) coding sequence file with default parameters. The mapping was done using kallisto quant for single-end library with an estimated fragment length of 200bp and estimated standard deviation of 20 option. In total, 41 experiments from *A. annua* were included in the co-expression network and annotated based on the information available from NCBI Sequence Read Archive. The Transcripts Per Kilobase Million (TPM) values from the kallisto outputs of selected experiments were represented as an expression matrix where genes were arranged in rows and experiments in columns (Table S1). To annotate the Artemisia CDS, we have used Mercator with standard settings [41]. Gene co-expression networks were calculated using Pearson Correlation Coefficient (PCC, [23]). The scripts used to perform these analyses are available from https://github.com/tqiaowen/LSTrAP-Lite.

## 3. RESULTS

### 3.1 Establishing the gene co-expression network pipeline on Rock64 single board computer

With the increased efficiency of transcript quantification softwares [35,38], estimating gene expression from RNA sequencing data is no longer a significant computational bottleneck. A recent blog post demonstrated that the kallisto program, used to estimate gene expression data, can successfully run on a credit card-sized single board computer Rock64 (https://liorpachter.wordpress.com/2018/01/29/bioinformatics-on-a-rock64/, Figure 1A).

We have established a computational pipeline that downloads the raw RNA-sequencing data from a public repository and constructs a gene expression matrix from the specified samples (Figure 1B). After compiling the expression matrix, the pipeline reports quality statistics of the individual samples by showing the number of reads that map to the coding sequences. This allows the user to remove samples that show poor mapping of the RNA-sequencing reads to the samples by specifying a list of selected accessions that should be used for downstream analyses. Finally, the pipeline accepts a gene of interest from the user to reveal co-expressed genes (Figure 1B). The co-expression relationships are reported as a co-expression list or a Cytoscape-compatible co-expression network [42].

To investigate the suitability of this single board computer in generating co-expression networks, we set to investigate the gene expression data of *Artemisia annua* [40], a plant that is cultivated globally as the main natural source of the potent anti-malarial compound, artemisinin. We used Rock64 to download and process all of the available RNA-seq accessions found in the European Nucleotide Archive and noted that most of the accession files are around 2 GB in size (Table S1). By using Aspera download client, we achieved average download speeds of 157 seconds per gigabyte (Figure 1C), with kallisto mapping taking 270 seconds per gigabyte. Overall, it took 4 hours to download and 7 hours to map 94 gigabytes of RNA-sequencing data (Table S1), showing that Rock64 is a viable platform to perform RNA-sequencing analyses. Analysis of the mapping statistics revealed one sample, SRR6472949 to map relatively poorly compared to the majority and was excluded from the analysis.

### 3.2 Elucidating the gene regulatory network for secondary cell wall biosynthesis

Secondary cell walls (SCW) provide mechanical support, water and nutrient transport and stress management in the vascular plant lineage. Since they also are an abundant resource of renewable feed, fiber, and fuel, a significant effort is directed to understand how SCWs are formed [43]. SCWs consist mainly of cellulose microfibrils and other polysaccharides such as hemicelluloses and pectins. In contrast to other cell wall types, SCWs contain lignin that makes the cell walls more rigid and less permeable to water [44]. Due to the importance of SCWs as a renewable resource, the biosynthetic and regulatory networks behind have received much attention and are well understood (Figure 2A). The transcription factors (TFs) controlling SCW are secondary wall NACs (SWNs), and are top-level master switches capable of inducing SCW formation ectopically [28]. The SWNs in turn regulate the expression and activity of several MYB transcription factors, that in turn activate expression of genes important for xylan, cellulose and lignin biosynthesis ([43], Figure 2A).

**Figure 2.**
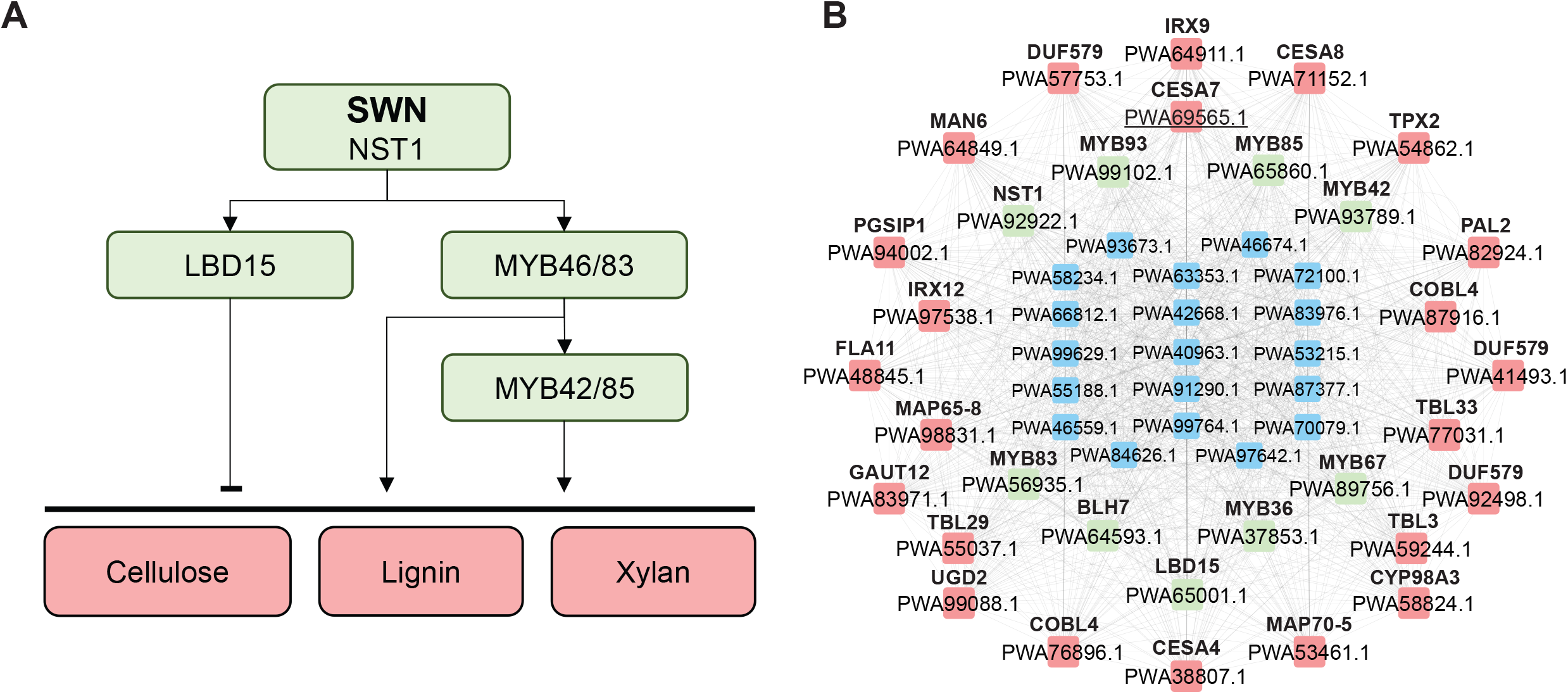
Regulatory and co-expression network of secondary cell wall biosynthesis in Artemisia. A) Schematic regulatory network of CESAs. Transcription factors are indicated in green, while their targets are shown in red. B) Co-expression network of CESA7 from Artemisia. Nodes represent genes, while edges connect co-expressed genes. Red nodes indicate genes involved in cell wall biosynthesis, green nodes indicate transcription factors, while blue nodes show other genes.

To demonstrate how co-expression networks can be used to reveal functionally related genes and GRN, we used our pipeline to retrieve 50 most co-expressed genes with *PWA69565.1*, a SCW-specific cellulose synthase 7 ortholog from *Artemisia* (Figure 2A, Table S3). The network revealed functionally related genes that are known participants of cellulose biosynthesis (CESAs [*PWA71152.1*, *PWA69565.1*, *PWA38807.1*], COBRAs [*PWA87916.1*, *PWA76896.1*] and TBL3 [*PWA59244.1*]) [45,46], deposition of cellulose microfibrils (FLA11 [*PWA48845.1*] and TPX2 [*PWA54862.1*]) [47,48], lignin biosynthesis (PAL2 [*PWA82924.1*], laccases [*PWA97538.1*], CYP450 [*PWA58824.1*]) [49,50], pectin biosynthesis (UGD2 [*PWA99088.1*]) [51], xylan biosynthesis (IRX9 [*PWA64911.1*], PGSIP1 [*PWA94002.1*], DUF579 [*PWA92498.1*, *PWA41493.1*, *PWA57753.1*], TBL29 [*PWA55037.1*], GAUT12 [*PWA83971.1*], TBL33 [*PWA77031.1*] [52–57] and cytoskeleton (MAPs [*PWA98831.1*, *PWA53461.1*]) [49].

Furthermore, several well studied TFs known for regulating various aspects of SCW formation were identified, such as NST1 (*PWA92922.1*), a master regulator of SCW biosynthesis; MYB83, MYB42 and MYB85 (*PWA56935.1*, *PWA93789.1* and *PWA65860.1* respectively) [43], positive regulators of SCW biosynthesis and LBD15 (*PWA65001.1*), a negative regulator of cellulose biosynthesis [58]. In addition to the conventional TFs associated with SCW, we also observe other TFs that may be involved in SCW in our network. For example, BLH7 was observed to be involved in the biosynthesis of cellulose-rich tension wood in *Populus* [59]. Suberin biosynthesis in the exocarp of cell wall of apples was proposed to be regulated by MYB93 [60]. Lastly, we also found MYB36, a TF associated with casparian strip formation, which consists of lignified primary cell walls [61].

Taken together, these results show that co-expression networks can find the biosynthetic genes and their regulators.

### 3.3 Elucidating the gene regulatory network for artemisinin biosynthesis

Malaria is one of the most deadly endemic infectious diseases. The World Health Organization reports that >1 billion people are at risk of malaria, and despite the recent efforts in the last decade, approximately 435,000 deaths were caused by malaria in 2017 [62,63]. Malaria is caused by microscopic parasites of the *Plasmodium* genus, among which *Plasmodium falciparum*, *P. vivax*, *P. ovale*, *P. malariae* and *P. knowlesi* are capable of infecting humans [64,65]. Unfortunately, *Plasmodium has* gained resistance to most medicines, such as quinine and chloroquine [64,66,67]. However, artemisinin from *Artemisia annua* is still an effective drug to treat malaria [68,69].

Artemisinin is an unusual endoperoxide sesquiterpene lactone [70] which is biosynthesized in the cytosol from isopentyl pyrophosphate (IPP) and dimethylallyl pyrophosphate (DMAPP, [71]). The biosynthetic network surrounding artemisinin also produces other valuable precursors, such as artemisinic acid and arteannuin B (Figure 3A, [72]). While the pathway is well understood and biosynthesis of artemisinin has been established in yeast [73], we hypothesize that other components, such as chaperones, co-factors or scaffolds could increase the biosynthesis of artemisinin [74].

**Figure 3.**
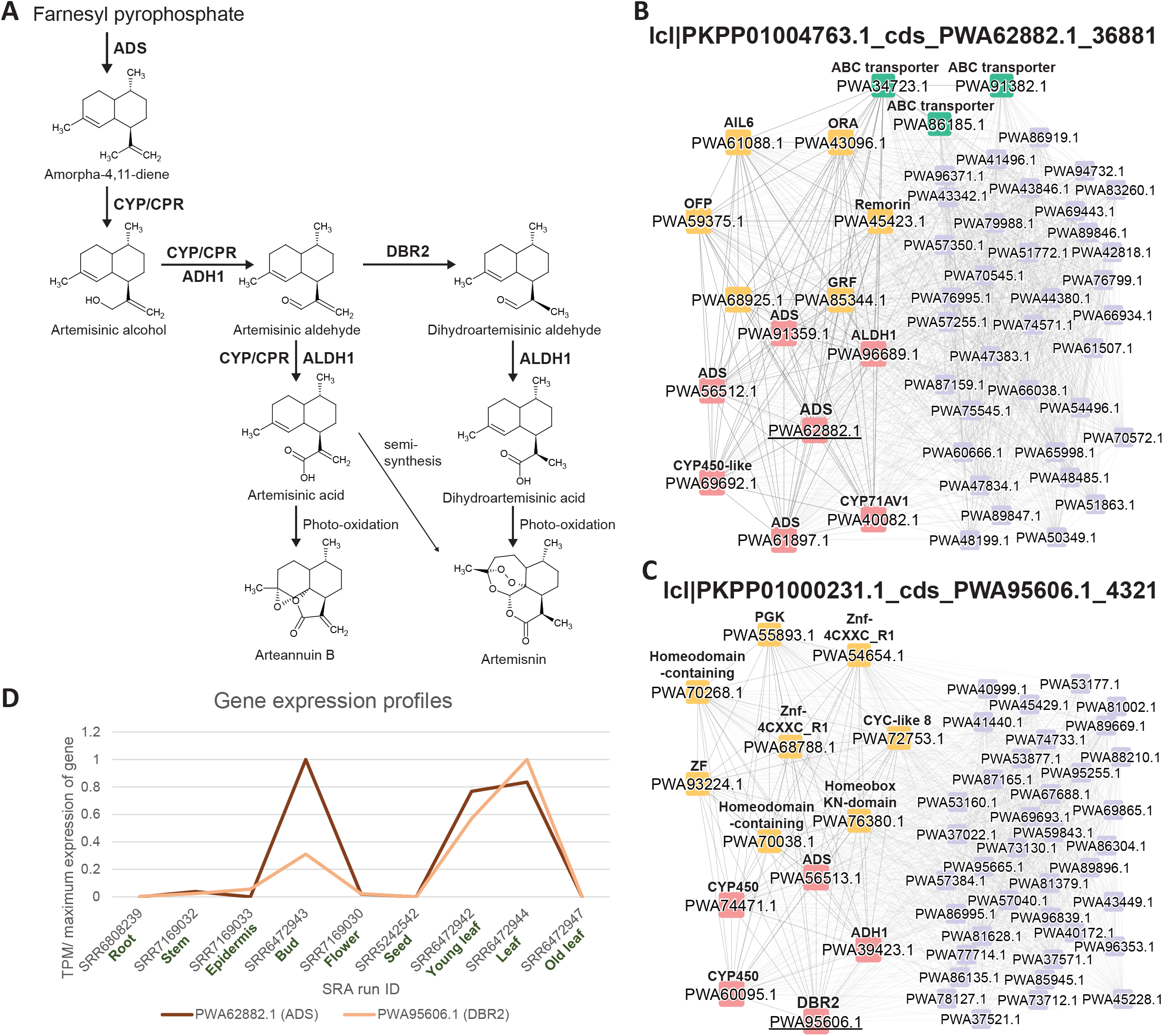
Expression profile and co-expression network of artemisinin biosynthesis pathway. A) A scheme showing biosynthesis of artemisinin. Abbreviations for enzymes include ADS, armorpha-4, 11-diene synthase, ADH1: alcohol dehydrogenase 1, ALDH1: aldehyde dehydrogenase, CPR: cytochrome P450 reductase, CYP71AV1: cytochrome P450 mono-oxygenase; DBR2: artemisinic aldehyde delta-11(13)-double bond reductase. B) Co-expression network of ADS (*PWA62882.1*). Red, orange and green nodes indicate enzymes, transcription factors and ABC transporters respectively, while blue nodes show other genes. Abbreviations for genes include AIL6: aintegumenta-like 6; GRF: growth regulating factor and OFP: ovate family protein. For brevity, only 50 most co-expressed genes of the query gene are shown. C) Co-expression network of DBR2 (*PWA95606.1*). Abbreviations for genes include Znf-4CXXC_R1: zinc-finger domain of monoamine-oxidase A repressor R1; CYC-like 8: cycloidea-like 8 and ZF: zinc finger. For brevity, only 50 most co-expressed genes of the query gene are shown. D) Expression profile of the two query genes from the shown co-expression networks. ADS and DBR2 are indicated by brown and orange colors, respectively.

To reveal the components involved in the biosynthesis of these high-value compounds, we used *PWA62882.1*, an amorpha-4,11-diene synthase (ADS) to retrieve its co-expression neighborhood (Figure 3B). The expression of *PWA62882.1* peaks in the bud and young leaf (Figure 3D). In the neighbourhood of *PWA62882.1*, we observe the presence of most genes involved in artemisinin biosynthesis [ADS (*PWA62882.1*, *PWA56512.1*, *PWA61897.1*), cytochrome P450-like (*PWA69692.1*), cytochrome P450 mono-oxygenase CYP71AV1 (*PWA40082.1*), alcohol dehydrogenase 1 (*PWA91359.1*) and aldehyde dehydrogenase 1 (*PWA96689.1*)] except for artemisinic aldehyde delta-11(13) reductase (DBR2) [75]. Notably, three ATP-binding cassette (ABC) transporters (*PWA91382.1*, *PWA86185.1*, *PWA34723.1*) are also found in the neighbourhood of *PWA62882.1*, making them interesting candidates for future optimisation of artemisinin biosynthesis as ABC transporters have been found to have a significant impact on artemisinin biosynthesis [76].

To date, various AP2/ERF, bHLH, WRKY and bZIP transcription factors have been reported to be involved in the positive regulation of artemisinin biosynthesis [76]. In the neighbourhood of *PWA62882.1*, we observe *PWA43096.1* (ORA) and *PWA68925.1*, a putative bZIP transcription factor, which may likely to play a role in the regulation of the artemisinin biosynthetic genes which co-expressed in the neighbourhood. Other transcription factors found in the network [aintegumenta-like 6 (*PWA61088.1*) [77], ovate family protein (*PWA59375.1*) [78], remorin (*PWA45423.1*) [79] and growth regulating factor (*PWA85344.1*) [80]] are not likely to be directly involved in the regulation of artemisinin biosynthesis but may be associated with their potential involvement in general growth and development of the plant under stress.

For a more complete picture of the genes and regulators involved in artemisinin biosynthesis, we looked at the co-expression neighbourhood of *PWA95606.1* (DBR2), the only gene in the biosynthetic pathway that was not identified in the neighbourhood of *PWA62882.1* (Figure 3C). The expression profile of *PWA95606.1* is similar to *PWA62882.1*, peaking at the bud and young leaf (Figure 3D). As expected, some genes involved in the biosynthesis of artemisinin were found in the neighbourhood of *PWA95606.1*, such as ADS (*PWA56513.1*), cytochrome P450 reductases (*PWA74471.1*, *PWA60095.1*) and alcohol dehydrogenase 1 (*PWA39423.1*). In contrast to the *PWA62882.1* neighbourhood, the DBR2 neighbourhood did not seem to associate with any of the well-studied transcription factors involved in the regulation of artemisinin biosynthesis. The transcription factors zinc-finger domain of monoamine-oxidase A repressor R1 (*PWA54654.1*, *PWA68788.1*), zinc finger (*PWA93224.1*), PGK (*PWA55893.1*), homeodomain-containing proteins (*PWA70268.1* and *PWA70038.1*), homeobox-KN domain containing protein (*PWA76380.1*) and cycloidea-like 8 (*PWA72753.1*) found in the neighbourhood of DBR2 also did not show any probable involvement in the regulation of artemisinin biosynthesis based on our current understanding. Thus, as exemplified in the examples described in Figure 3B and 3C, the choice of gene to be used as bait is highly crucial and can sometimes give very different or incomplete perspectives on the biological process of interest.

## 4. DISCUSSION

Gene function prediction and inference of GRN are in the focal point of bioinformatics, and have been the subject of numerous DREAM challenges, which invite participants to compete in providing the most accurate and complete predictions [30,81]. The conclusions from these challenges and other studies is that integration of multiple data types and inference methods typically improves performance of the bioinformatical predictions [1,82–84].

Here, we show that a simple co-expression analysis is able to infer regulators, enzymes and structural proteins involved in secondary cell wall synthesis and artemisinin in *Artemisia* (Figure 2–3). These results show that biosynthetic pathways and their regulators can be readily inferred by this routine analysis [33,85–88]. However, it remains unclear whether there is a correlation between the predictability of biosynthetic pathways of secondary metabolism and predictability of the regulatory networks controlling them. Secondary metabolism genes tend to have higher proliferation rates by local gene duplications, are often co-localized in the genome and are co-expressed [89], but these observations cannot explain why these genes are often co-expressed with transcription factors controlling them. While these findings suggest that, in contrast to other processes, secondary metabolism is perhaps under simpler, exclusively transcriptional control, further work is needed to confirm this hypothesis.

Affordable computing and availability of biological data is providing us with tools to generate high quality predictions of gene function and GRNs. The ease and affordability of generating and analyzing new data requires an overhaul of education and training of all researchers, from undergraduates to faculty [90]. These steps are already seen across the globe, with topics such as computational thinking, programming and data science being introduced as compulsory subjects for biology undergraduates.

## DATA AVAILABILITY

The expression matrix is found in the supplemental data, while the scripts used on Rock64 are found at https://github.com/tqiaowen/LSTrAP-Lite

## SUPPLEMENTARY DATA

**Table S1. Sample annotation, download times and kallisto mapping statistics for Artemisia.**

**Table S2. Gene annotation and expression matrix of *Artemisia attenuata***. The first, second and third columns contain gene identifiers, gene annotations from Mercator and MapMan identifiers, respectively. The expression profiles start from the 4th column.

**Table S3 Table of top 50 co-expressed genes with the *PWA68565.1* (*CESA7*).**

**Table S4. Table of top 50 co-expressed genes with *PWA62882.1* (*ADS*).**

**Table S5. Table of top 50 co-expressed genes with *PWA95606.1* (*DBR2*).**

## ACKNOWLEDGEMENT

We would like to thank Dr. Daniela Mutwil-Anderwald for constructive comments on this manuscript.

